# Sequential gentle hydration increases encapsulation in model protocells

**DOI:** 10.1101/2023.10.15.562404

**Authors:** Emma M. Gehlbach, Abbey O. Robinson, Aaron E. Engelhart, Katarzyna P. Adamala

## Abstract

Small, spherical vesicles are a widely used chassis for the formation of model protocells and investigating the beginning of compartmentalized evolution. Various methods exist for their preparation, with one of the most common approaches being gentle hydration, where thin layers of lipids are hydrated with aqueous solutions and gently agitated to form vesicles. An important benefit to gentle hydration is that the method produces vesicles without introducing any organic contaminants, such as mineral oil, into the lipid bilayer. However, compared to other methods of liposome formation, gentle hydration is much less efficient at encapsulating aqueous cargo. Improving the encapsulation efficiency of gentle hydration would be of broad use for medicine, biotechnology, and protocell research. Here, we describe a method of sequentially hydrating lipid thin films to increase encapsulation efficiency. We demonstrate that sequential gentle hydration significantly improves encapsulation of water-soluble cargo compared to the traditional method, and that this improved efficiency is dependent on buffer composition. Similarly, we also demonstrate how this method can be used to increase concentrations of oleic acid, a fatty acid commonly used in origins of life research, to improve the formation of vesicles in aqueous buffer.

## Introduction

The most common vessel used in biology, and a tool for investigation of the origins and earliest evolution of compartmentalized life, are liposomes, also called unilamellar vesicles (UVs) (Bogdan-Cezar and Stano, 2023; Smoukov et al., 2023; Subbotin and Fiksel, 2023). Compartmentalization was likely essential for the emergence of life, as all biological cells are bound by a plasma membrane (Jin et al., 2018; Monnard and Walde, 2015). Liposomes consist of a spherical lipid bilayer containing an aqueous inner solution (the lumen). They can act as a simple model of cell membranes, making them an indispensable tool in studies of the origin of biological encapsulation. Liposomes can be cell-sized, ranging from around 0.1 to 100 microns in diameter, and they often are constructed from phospholipids and cholesterol (Akbarzadeh et al., 2013). These cell-like microcontainers have applications beyond the studies of the origins of life in synthetic biology, drug delivery, cosmetics, and as bioreactors (Akbarzadeh et al., 2013; Daraee et al., 2014; Dewey et al., 2014; Rahimpour and Hamishehkar, 2012; Sato et al., 2022). There are various methods of encapsulating aqueous cargo in liposomes, and these approaches can be defined as either active or passive (Hood et al., 2014). Active loading utilizes chemical, ionic, or pH gradients to encapsulate compounds in already assembled liposomes. Active loading has high encapsulation efficiency; however, this efficiency is dependent on the lipid composition and quality, solubility and pKa of the compound, the presence of functional groups on the compound, and other factors (Haran et al., 1993; Sur et al., 2014). In the passive loading process, the internal solution is added to the lipids prior to the formation of vesicles, and liposomes are formed as lipids assemble around the aqueous droplets. Examples of passive loading include gentle hydration (also called natural swelling), electroformation, and water-in-oil emulsion (Akbarzadeh et al., 2013; Angelova and Dimitrov, 1986; Osaki et al., 2011; Venero et al., 2022). During gentle hydration, aqueous buffer is added to dried lipid thin films, and agitated by a combination of sonication, vortexing, and rotation resulting in the “swelling” of vesicles containing the buffer (Tsumoto et al., 2009). However, passive loading tends to have lower encapsulation efficiencies compared to active loading, which can result in wasted compounds.

Of the passive loading techniques, water-in-oil emulsion has the highest encapsulation efficiency of up to 65% (Akbarzadeh et al., 2013). However, this method often results in residual oil left between the leaflets of the membrane of liposomes (Kamiya et al., 2016). The presence of residual solvent creates an uncontrolled variable during experiments and can negatively impact experimental results if not accounted for. Residual solvent has been shown to affect the rigidity and fluidity of the membrane, which can in turn affect membrane dynamic and protein behavior (Booth et al., 1997; Lorch and Booth, 2004; van Swaay and deMello, 2013). Thus, there is a need to increase encapsulation efficiencies of passive loading techniques that do not utilize mineral oil or organic solvent. Here, we describe a method of sequential gentle hydration to increase encapsulation efficiency, while introducing no mineral oil to the lipid mixture. We hydrated lipid thin films in aqueous buffer containing a fluorescent dye, to create liposomes, and the buffer containing the liposomes and un-encapsulated dye are then added to hydrate a series of thin films to increase encapsulation of dye. We tested this method using two common buffers, Tris and HEPES. Using this method with HEPES buffer, we observed nearly a six-fold increase in encapsulation compared to traditional gentle hydration. Buffers containing HEPES are used in transcription-translation (TXTL) systems, recreating model protocells and the last universal common ancestor (LUCA) (Gaut and Adamala, 2021; Mann, 2013). The ability to improve encapsulation within lipid vesicles through sequential gentle hydration in HEPES buffer indicates that sequential hydration can be a viable method to improve the yield for protocells, and for other research utilizing cell-free and other enzymatic systems (Sun et al., 2013). We also use this method to increase the encapsulation of cargo in vesicles made from oleic acid, the fatty acid predominantly used in origins of life research. We used bicine buffer when making oleic acid vesicles, as it is commonly used with those lipids.

## Materials and Methods

### POPC and cholesterol thin film preparation

Thin films were prepared by dissolving 1-palmitoyl-2-oleoyl-sn-glycero-3-PC (POPC) and cholesterol (Cayman Chemical) in chloroform to create 50 mg/mL solutions that were partitioned into amber glass vials. Vials were prepared containing 1 mM POPC and 1 mM cholesterol concentration for sequential gentle hydration, and vials containing 1-9 mM POPC/cholesterol were prepared for non-sequential gentle hydration. Vials were left uncapped overnight in a fume hood to allow for the solvent to evaporate and thin films to form. Thin films were capped and stored at -20°C.

### Oleic acid thin film preparation

To prepare thin films, oleic acid (Cayman Chemical) was dissolved in chloroform to assist in pipetting the viscous liquid. The solution was partitioned into amber glass vials, such that thin films for sequential gentle hydration contained 5 mM oleic acid and thin films for non-sequential gentle hydration contained 10, 15, and 20 mM oleic acid. Vials were left uncapped in the fume hood to allow the chloroform to evaporate overnight. The vials containing oleic acid were capped and stored at room temperature.

### Sequential gentle hydration

1 mL of either CHAN buffer (10 mM Tris, 100 mM NaCl, 1 mM disodium-EDTA, pH 7.0), HEPES buffer (100 mM HEPES, pH 7.6), or bicine buffer (100 mM bicine, pH 8.0) containing 0.5 mM calcein disodium dye was added to lipid thin films. Liposomes were formed through gentle hydration by vortexing each aliquot for 1 minute, sonicating for 15 minutes at 60°C, vortexing for another minute, sonicating for another 15 minutes at 60°C, and vortexing for 1 final minute. Vials were put on a rotator and were left on a rotator for 12-48 hours. For the sequential process, lipids were incorporated into the solution by adding 1 mM POPC and 1 mM cholesterol or 5 mM oleic acid at a time and vortexing and sonicating accordingly. After each cycle of lipid addition and vortexing/sonication, aliquots were put on a rotator and were left on the rotator for 12-48 hours, until the next step of the process. No more than one addition of lipids was performed per day, and vials containing oleic acid were left on the rotator for at least 24 hours after each lipid addition. For the non-sequential control process, all the lipids were resuspended from one thin film and the vortexing and sonication process was performed once. The final concentrations of cholesterol and POPC for both methods increased stepwise in increments of 1 mM of both POPC and cholesterol. The final concentrations of oleic acid in solution increased stepwise in increments of 5 mM.

### Size extrusion

Liposomes were extruded using a Mini-Extruder (Avanti), and filters and drain discs (Whatman). The extruder was assembled using a 19 mm 1.0-μm Nucleopore Track-Etched membrane extrusion filter and a 10 mm PE drain disc. The Teflon bearing included in the Mini-Extruder assembly was not used. The liposome-containing solutions were pushed through the filter 7 times between two 1-mL gas-tight glass syringes (Hamilton) to remove liposomes with a diameter larger than 1 μm from the sample. A small loss of sample volume occurred during the extrusion process.

### Sepharose purification and measuring encapsulation

Size-exclusion chromatography (SEC) was performed to separate the liposomes from the unencapsulated dye in the buffer. A 10 mL column containing Sepharose 4B (Sigma Aldrich) was washed 4 times with buffer before liposomes were loaded. 150 μL of each extruded liposome sample was gently loaded onto the Sepharose and buffer was used to wash the liposomes through the column. The eluate was fractionated into a 96-well non-treated round bottom plate (CellTreat). The plates were read on a microplate fluorometer (SpectraMax GeminiXS) to measure calcein fluorescence (excitation: 495 nm; emission: 515 nm; readings: precise (30); PMT: low) and the fluorescence values at each well were recorded. The percentage of the buffer solution that was encapsulated within the liposomes was calculated using Formula 1. The column was washed with at least one volume of buffer between different samples.

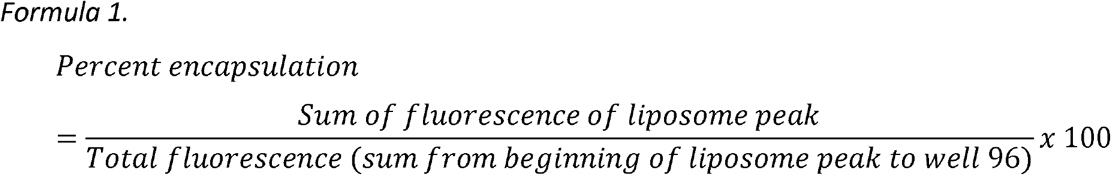

## Results

### Sequential gentle hydration increases encapsulation efficiency in HEPES buffer

For sequential gentle hydration, 0.5 mM calcein disodium dye was added to HEPES liposome buffer for the internal solution. 1 mL of the internal solution was added to 1 mM POPC/cholesterol thin films in glass vials. The vials were then vortexed, sonicated, and put on a rotator overnight for formation of liposomes. The next day, the solution containing the 1 mM POPC/cholesterol liposomes and unencapsulated buffer was added to a new 1 mM thin film. The vial was then vortexed, sonicated, rotated overnight, and this process was repeated to create up to a 9 mM lipid mixture. When the lipid solutions reached their final concentration, they were extruded through a 1 μm filter to reduce size dispersion of the liposomes for size exclusion chromatography (SEC).

To understand the degree of which gentle hydration of each additional thin film increased encapsulation efficiency, SEC was used to separate liposomes encapsulating calcein from the free calcein within the buffer. During SEC, the gentle hydration mixture is loaded into a column containing Sepharose gel beads with large pores that trap solute particles that are not encapsulated, while the larger liposomes are able to pass through the gel beads and are eluted first from the column (Grabielle-Madelmont et al., 2003). The samples were loaded onto a column containing Sepharose 4B, fractionated into a microplate, and fluorescence of the fractions was measured on a plate reader. The extruded liposomes were eluted first in the microplate wells, followed later by the unencapsulated buffer. Thus, two distinct fluorescent peaks were observed: the peak from the first wells on the microplate represent the fluorescence signal from the calcein trapped in liposomes that exit the column first, and the second peak from the later fractions represents the calcein dissolved in the unencapsulated buffer. The sum of the area under the peaks was used to determine the percent encapsulation (Fig. 1). The percent encapsulation was compared between liposomes prepared by sequential gentle hydration and non-sequential gentle hydration.

**Figure 1.**
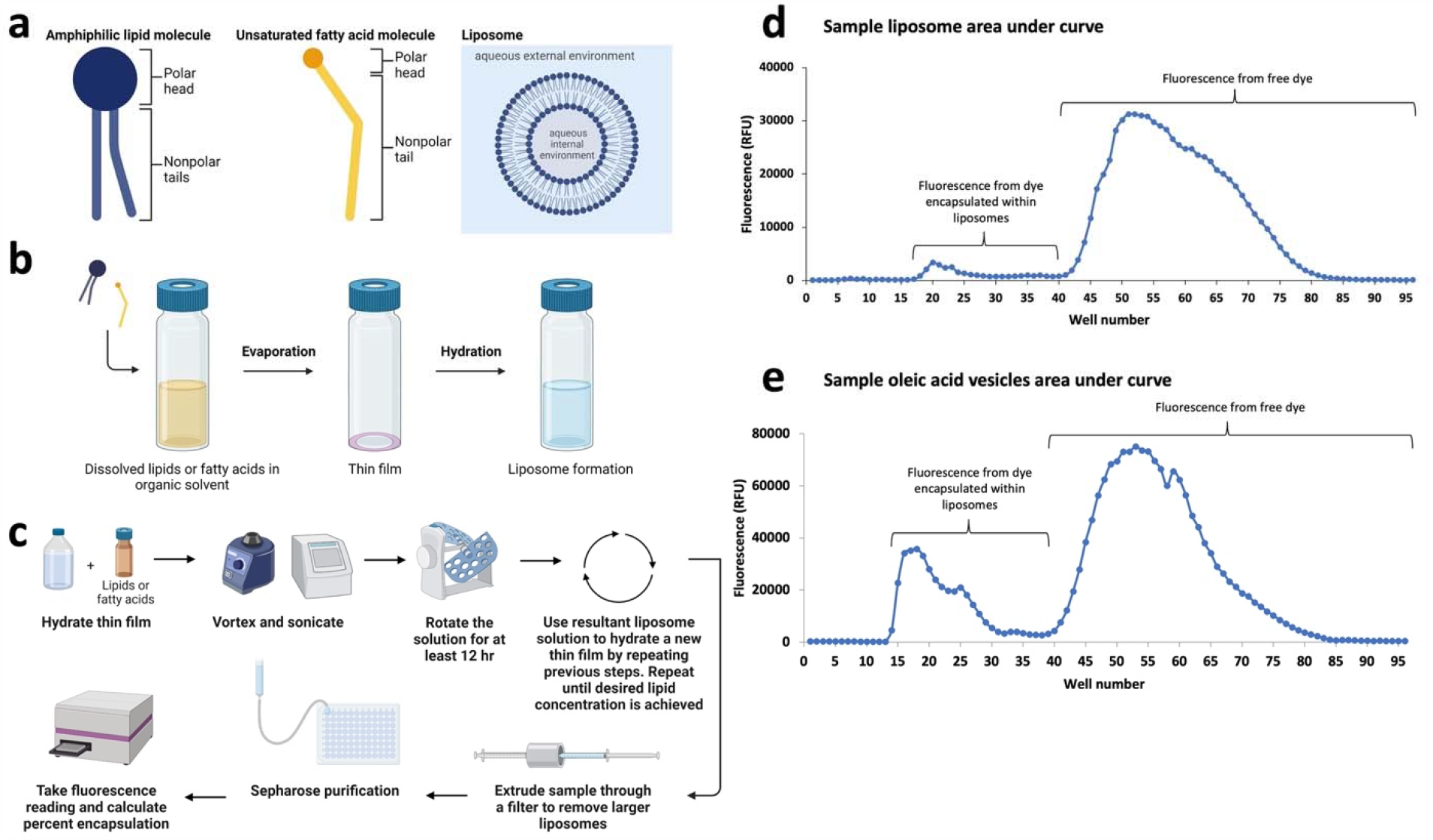
Overview of sequential gentle hydration method. a) Schematic of lipid and fatty acid monomers and liposomes. b) Schematic of thin film preparation and hydration. c) Workflow for sequential gentle hydration method. Thin films are hydrated with aqueous buffer, vortexed and sonicated, and put on a rotator for 12-48 hours. The resultant solution containing liposomes and residual unencapsulated material are added to a new thin film and the process is repeated until the desired lipid concentration is achieved. Liposomes are extruded through a filter and purified using Sepharose size exclusion chromatography (SEC). Fluorescence reads are taken, and the percent encapsulation is calculated. d) Example fluorescence data used to calculate percent encapsulation for POPC/cholesterol liposomes. During SEC, the purified liposomes and unencapsulated dye are fractionated onto a 96-well microplate. The plate is then read using a fluorometer. The resulting figure depicts two peaks correlating to the fractions: the first, smaller peak reflects the dye encapsulated in vesicles which is eluted first; the second, larger peak reflects the unencapsulated dye eluted last. e) Example fluorescence data for oleic acid used to calculate percent encapsulation.

Liposomes prepared by traditional gentle hydration had between 0.5-1.5% average encapsulation for thin films ranging from 1-9 mM lipid concentration. The sequential gentle hydration method achieved an average of 5.7% encapsulation at 8 mM lipid concentration, an almost 6-fold increase in encapsulation compared to the traditional method at 8 mM concentration. Sequential gentle hydration was able to produce significantly higher percent encapsulation compared to non-sequential hydration at 4, 5, 6, and 7 mM POPC and cholesterol. However, there is a high amount of variability in encapsulation when using the sequential gentle hydration method, which meant that the highest concentrations of sequentially hydrated POPC and cholesterol (8 and 9 mM) did not exhibit significantly higher encapsulation efficiency than the non-sequentially hydrated lipids at the same concentrations. Interestingly, the average encapsulation efficiency fell off between 8 mM (5.7%) and 9 mM (4.3%) (Fig. 2a-b, Table 1). We hypothesize this high variability could be due to the longer rotation times needed to sequentially encapsulate higher lipid concentrations, which may repeatedly jostle and disturb the liposomes, causing them to rupture. This may have freed lipid monomers, increasing the concentration of available lipids in the solution such that more lipid monomers can interact with each other, allowing multilamellar or larger liposomes to form.

**Table 1.** Percent encapsulation of calcein disodium dye in POPC/cholesterol vesicles. Comparison between sequential and non-sequential gentle hydration in HEPES- and Tris-based buffers. *\* p ≤ 0*.*05, ** p ≤ 0*.*01*.

| POPC and cholesterol concentration (mM) | Average % encapsulation in HEPES buffer, sequential | Average % encapsulation in HEPES buffer, non-sequential | Average % encapsulation in Tris buffer, sequential | Average % encapsulation in Tris buffer, non-sequential |
| --- | --- | --- | --- | --- |
| 1<sup>†</sup> | 0.55<sup>†</sup> | 0.55<sup>†</sup> | 0.93<sup>†</sup> | 0.93<sup>†</sup> |
| 2 | 0.68 | 0.56 | 1.06 | 1.05 |
| 3 | 0.80 | 0.76 | 1.50 | 0.99 |
| 4 | 1.96* | 0.81 | 1.34 | 1.43 |
| 5 | 2.25** | 0.93 | 1.20 | 1.34 |
| 6 | 2.40** | 1.44 | 1.18 | 1.84* |
| 7 | 4.00* | 1.12 | 1.09 | 1.49 |
| 8 | 5.72 | 0.96 | 1.68 | 1.25 |
| 9 | 4.25 | 1.22 | 1.59 | 1.60 |

<sup>†</sup> Percent encapsulation for the starting lipid concentration is a reference value for both data sets respective to buffer, since only one thin film was used.

**Figure 2.**
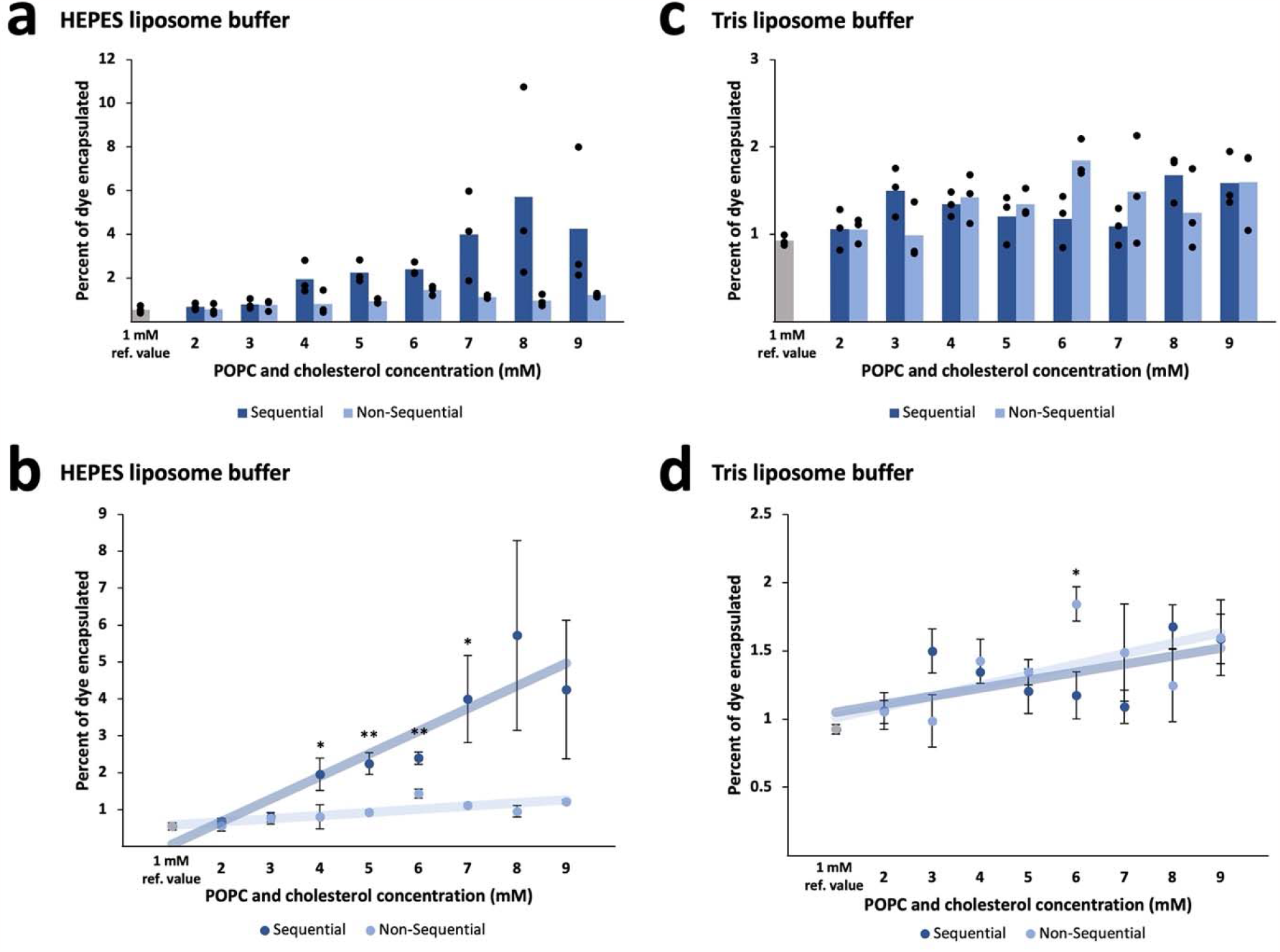
Percent encapsulation of calcein disodium dye in POPC/cholesterol liposomes made by sequential gentle hydration in HEPES and Tris buffer. a) Fluorescence measurements for liposomes made by the sequential and traditional methods were taken in triplicate for each lipid concentration. The sequential and non-sequential processes for HEPES buffer are shown with a linear regression to demonstrate the relationship between lipid concentration and percent encapsulation. The 1 mM samples were included for reference to typical percent encapsulation of calcein in HEPES buffer, and they were included in both the sequential and non-sequential categories when calculating the trendlines. b) The average percent encapsulation of calcein within liposomes in HEPES buffer is shown for both the sequential and non-sequential processes at each lipid concentration. For HEPES buffer, the sequential method results in higher percent encapsulation compared to the non-sequential method. c) The sequential and non-sequential processes for Tris liposome buffer. d) The average percent encapsulation of calcein within liposomes in Tris liposome buffer is shown for both the sequential and non-sequential processes at each lipid concentration. The sequential method was not as effective in the Tris liposome buffer. *\* p ≤ 0*.*05, ** p ≤ 0*.*01*.

### Effect of buffer composition on encapsulation efficiency

The effect of buffer composition on encapsulation efficiency was also tested. Sequential gentle hydration was done with two common buffering agents, Tris and HEPES. Tris and HEPES are both suitable for liposome preparation and storage. Notably, HEPES is used in TXTL protein expression systems. For the Tris-containing buffer, we used our standard liposome storage buffer which contains 10 mM Tris, 100 mM NaCl, and 1 mM disodium-EDTA.

Interestingly, the encapsulation rate for sequential gentle hydration with the Tris-based liposome buffer was considerably lower compared to when HEPES buffer was used. There was no notable increase in encapsulation between the sequential method and traditional method when the Tris liposome buffer was used (Fig. 2c-d, Table 1). We hypothesize this is due to the presence of additional salt in the buffer, which is known to suppress vesicle formation as Na^+^ and Cl^−^ ions interact with the lipid heads. Stronger interactions between lipid heads and the buffer/salts leads to stronger sequestration of lipid tails away from the aqueous environment, causing tighter packing of lipid molecules and reduced lateral movement. This inhibits liposome formation and promotes formation of micelles, which are not capable of encapsulating aqueous solutions. This likely explains why encapsulation efficiency was reduced in the Tris buffer that contained 100 mM NaCl (Wang et al., 2017).

### Sequential gentle hydration of oleic acid

We also wished to explore the effect of membrane composition on encapsulation efficiency using the sequential hydration method. We tested our method on vesicles made from oleic acid, a fatty acid that forms ufasomes (unsaturated fatty acid vesicles). Fatty acids are simpler, more stable, and more permeable than phospholipids, and oleate is often used to model prebiotic membranes. Phospholipids and sterols are biosynthesized by specialized enzymatic pathways that were not yet in existence at the time of the creation of the first cellular compartments; thus, simpler molecules are used to study early membranes, and their high stability allows them to accumulate under unfavorable conditions even when they are not being actively synthesized (Deamer, 2017; Schrum et al., 2010). Oleic acid-based vesicles are also used for drug delivery (Kumar et al., 2018; Zakir et al., 2010).

For sequential gentle hydration, bicine buffer (100 mM, pH 8.0) containing 0.5 mM calcein dye was used to hydrate oleic acid thin films. The thin films were dissolved sequentially in 5 mM increments as described in Methods. Ufasomes made by sequential gentle hydration nearly doubled the encapsulation efficiency compared to ufasomes made through traditional, non-sequential gentle hydration. Notably, when thin films were sequentially dissolved over a period of days, higher concentrations of oleic acid (25-35 mM) were able to be dissolved in the buffer, whereas these higher concentrations of oleate did not fully dissolve during the traditional overnight non-sequential hydration. These higher concentrations of oleic acid produced higher encapsulation efficiencies up to 17.3%, which was significantly larger than the highest average encapsulation efficiency of the non-sequential samples (5.99%) (Fig. 3, Table 2). Additionally, higher concentrations of non-sequentially hydrated oleic acid were more difficult to extrude than sequentially hydrated oleic acid at the same concentration, suggesting that non-sequential process tends to form a greater number of large ufasomes that increase the force needed to extrude the vesicle-containing solution through the 1 μm filter. Improving the encapsulation efficiency of oleic acid ufasomes will be beneficial for applications in biotechnology as well as research into the origins of life. For example, it has been proposed that channels in minerals near hydrothermal vents provided compartments where biological molecules slowly accumulated and reached high enough concentrations to form liposomes that would eventually become the first protocells (Martin and Russell, 2003; Schrum et al., 2010). The process of sequentially hydrating lipids can mimic the slow increase in lipid concentration in these environments over time more effectively than non-sequential hydration.

**Table 2.** Percent encapsulation of calcein disodium dye in oleic acid vesicles. Comparison between sequential and non-sequential gentle hydration. *\* p ≤ 0*.*05, ** p ≤ 0*.*01*.

| Oleic acid concentration (mM) | Average % encapsulation in bicine buffer, sequential | Average % encapsulation in bicine buffer, non-sequential |
| --- | --- | --- |
| 5<sup>††</sup> | 2.64<sup>††</sup> | 2.64<sup>††</sup> |
| 10 | 5.81 | 3.89 |
| 15 | 9.90* | 5.57 |
| 20 | 11.15 | 5.99 |
| 25 | 12.53 |  |
| 30 | 17.35 |  |
| 35 | 16.82 |  |

<sup>††</sup> Percent encapsulation for the starting oleic acid concentration is a reference value for both data sets, since only one thin film was used.

**Figure 3.**
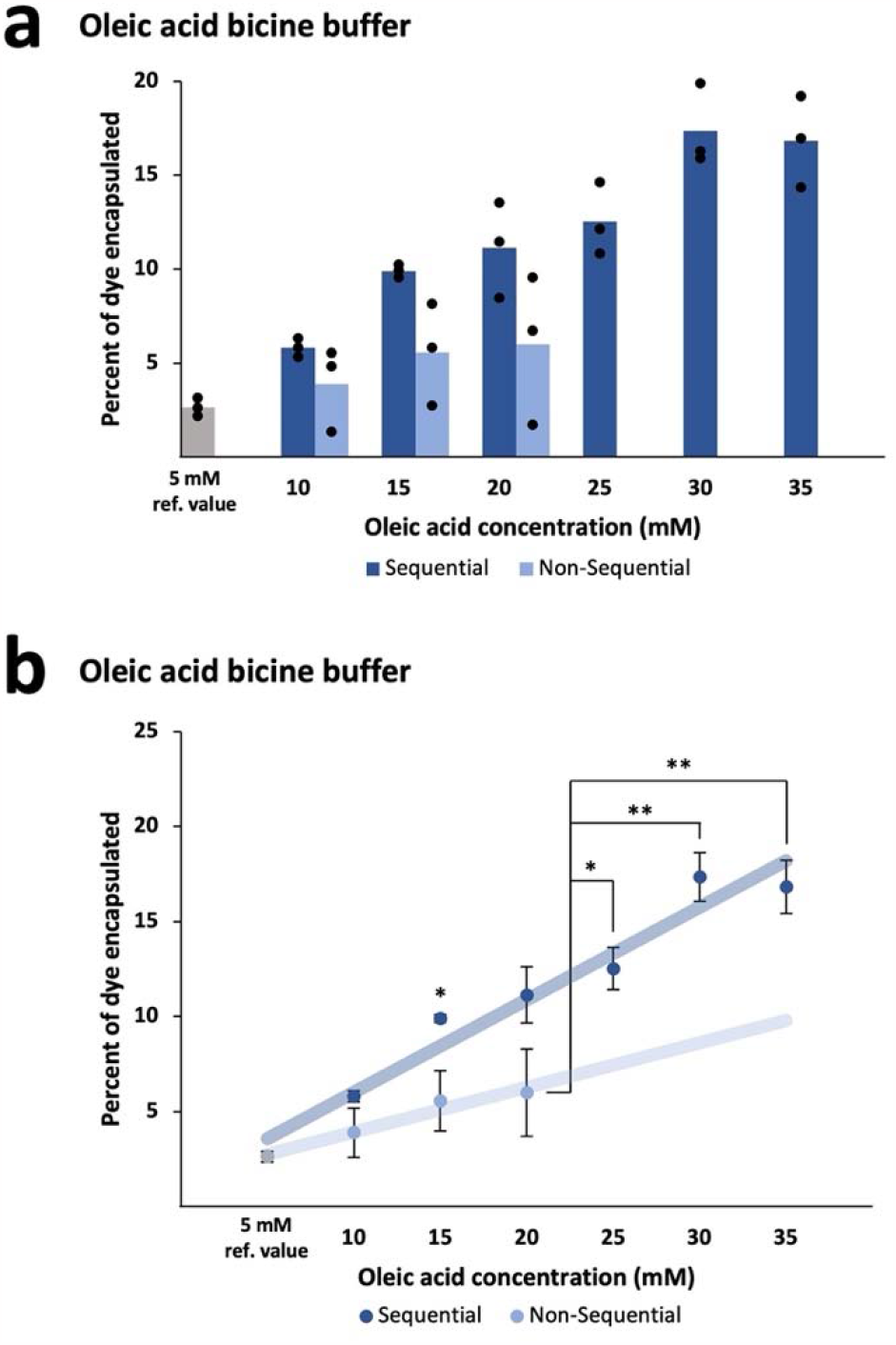
Percent encapsulation of calcein disodium dye in oleic acid vesicles made by sequential gentle hydration in bicine buffer. a) Fluorescence measurements for ufasomes made by the sequential and traditional methods were taken in triplicate for each oleic acid concentration. The sequential and non-sequential processes are shown with a linear regression to illustrate the relationship between oleic acid concentration and encapsulation efficiency. The 5 mM samples were included for reference to typical encapsulation efficiency of calcein in bicine buffer and were included in both the sequential and non-sequential categories when calculating the trendlines. b) The average percent encapsulation of calcein in liposomes is shown for both the sequential and traditional methods at each oleic acid concentration. The sequential method produced higher encapsulation efficiency than the sequential method and increased the dissolvable concentration of oleic acid to form higher-concentration oleic acid solutions that produced higher percent encapsulation values than the traditional method. *\* p ≤ 0.05, ** p ≤ 0.01*.

## Conclusions

Sequential gentle hydration is an effective method for improving the encapsulation efficiency of aqueous cargo by passive loading. Sequential gentle hydration of POPC/cholesterol thin films in HEPES buffer and oleic acid thin films in bicine buffer improved encapsulation rates for water-soluble small molecules, compared to non-sequential gentle hydration. While there was increased encapsulation using this method, we noted a wide distribution of encapsulation efficiencies using the sequential method. This wide distribution may have occurred due to experimental constraints (the time needed to dissolve the lipids), or it could be an inherent property of high concentration lipid systems. We found that buffer composition influences the formation of vesicles using the sequential method: while the method was effective using HEPES, the sequential process did not produce significantly higher average percent encapsulation in the Tris-based buffer. Notably, that buffer contains 100 mM NaCl, and liposome formation is reduced in salt solutions with a concentration higher than 50 mM (Wang et al., 2017). Encapsulation of dye within oleic acid vesicles made by sequential gentle hydration nearly doubled compared to ufasomes made through the non-sequential method, and higher concentrations of oleic acid were possible only when the sequential method was used. Those observations might provide insights for designing high salt liposome experiments, under conditions simulating one of the possible prebiotic environments (Damer and Deamer, 2020).

Improving encapsulation in the gentle hydration process through sequential gentle hydration in HEPES, which is the buffer used in protein expression and other enzymatic reactions, expands the usefulness of gentle hydration in model protocell research. In some scenarios, gentle hydration may be able to replace the emulsion method of forming liposomes, yielding protocells that are more similar to biological cells, and therefore, these liposomes can more accurately model LUCA and its immediate ancestors (Adamala and Luisi, 2011; Osaki et al., 2011; Stano et al., 2012). Expanding use of gentle hydration would ensure that the liposomes do not have residual organic solvent in the bilayer, removing an uncontrolled variable from the experiment (Booth et al., 1997; Kamiya et al., 2016; Lorch and Booth, 2004; Osaki et al., 2011; van Swaay and deMello, 2013). Additionally, sequential gentle hydration increased the encapsulation efficiency, and the dissolvable concentration of oleic acid forming ufasomes. This improved method of forming oleic acid protocells can be used in many areas of origins of life research (Deamer, 2017; Martin and Russell, 2003; Schrum et al., 2010). Sequential gentle hydration may be used to investigate the theory that channels in minerals near hydrothermal vents provided compartments where biological molecules slowly accumulated, reaching high enough concentrations to form protocell liposomes (Martin and Russell, 2003; Schrum et al., 2010). Sequential hydration is a realistic model for this process because it can mimic the slow increase in lipid concentration in these environments over time. Overall, we present a tool for foundational astrobiology research, and with potential utility for synthetic biology and biomedical applications.

## Acknowledgements

Figures were created using BioRender.com.

## Authorship Statement

*Emma M. Gehlbach*: Investigation, Formal Analysis, Visualization, Writing – Original Draft, *Abbey O. Robinson*: Investigation, Supervision, Visualization, Writing – Original Draft, Project Administration, *Aaron E. Engelhart*: Funding acquisition, Methodology, Writing – Review & Editing, Resources, *Katarzyna P. Adamala*: Conceptualization, Methodology, Supervision, Resources, Visualization, Writing – Review & Editing, Project Administration, Funding acquisition

## References

Adamala K and Luisi PL. Experimental Systems to Explore Life Origin: Perspectives for Understanding Primitive Mechanisms of Cell Division. In: Cell Cycle in Development. (Kubiak JZ. ed) Springer Berlin: Heidelberg; 2011; pp. 1–9; doi: 10.1007/978-3-642-19065-0_1.

Akbarzadeh A, Rezaei-Sadabady R, Davaran S, et al. Liposome: Classification, Preparation, and Applications. Nanoscale Res Lett 2013;8(1):102; doi: 10.1186/1556-276X-8-102.

Angelova MI and Dimitrov DS. Liposome Electroformation. Faraday Discuss Chem Soc 1986;81:303–311; doi: 10.1039/dc9868100303.

Bogdan-Cezar I and Stano P. Chemical Systems for Wetware Artificial Life: Selected Perspectives in Synthetic Cell Research. International Journal of Molecular Sciences 2023, Vol 24, Page 14138 2023;24(18):14138; doi: 10.3390/IJMS241814138.

Booth PJ, Riley ML, Flitsch SL, et al. Evidence That Bilayer Bending Rigidity Affects Membrane Protein Folding. Biochemistry 1997;36(1):197–203; doi: 10.1021/bi962200m.

Damer B and Deamer D. The Hot Spring Hypothesis for an Origin of Life. Astrobiology 2020;20(4):429–452; doi: 10.1089/ast.2019.2045.

Daraee H, Etemadi A, Kouhi M, et al. Application of Liposomes in Medicine and Drug Delivery. Artif Cells Nanomed Biotechnol 2014;44(1):381–391; doi: 10.3109/21691401.2014.953633.

Deamer D. The Role of Lipid Membranes in Life’s Origin. Life 2017;7(1):5; doi: 10.3390/life7010005.

Dewey DC, Strulson CA, Cacace DN, et al. Bioreactor Droplets from Liposome-Stabilized All-Aqueous Emulsions. Nat Commun 2014;5(1):4670; doi: 10.1038/ncomms5670.

Gaut NJ and Adamala KP. Reconstituting Natural Cell Elements in Synthetic Cells. Adv Biol 2021;5(3):e2000188; doi: 10.1002/adbi.202000188.

Grabielle-Madelmont C, Lesieur S and Ollivon M. Characterization of Loaded Liposomes by Size Exclusion Chromatography. J Biochem Biophys Methods 2003;56(1–3):189–217; doi: 10.1016/S0165-022X(03)00059-9.

Haran G, Cohen R, Bar LK, et al. Transmembrane Ammonium Sulfate Gradients in Liposomes Produce Efficient and Stable Entrapment of Amphipathic Weak Bases. Biochimica et Biophysica Acta (BBA) - Biomembranes 1993;1151(2):201–215; doi: 10.1016/0005-2736(93)90105-9.

Hood RR, Vreeland WN and DeVoe DL. Microfluidic Remote Loading for Rapid Single-Step Liposomal Drug Preparation. Lab Chip 2014;14(17):3359–3367; doi: 10.1039/C4LC00390J.

Jin L, Engelhart AE, Adamala KP, et al. Preparation, Purification, and Use of Fatty Acid-Containing Liposomes. Journal of Visualized Experiments 2018;(132); doi: 10.3791/57324.

Kamiya K, Kawano R, Osaki T, et al. Cell-Sized Asymmetric Lipid Vesicles Facilitate the Investigation of Asymmetric Membranes. Nat Chem 2016;8(9):881–889; doi: 10.1038/nchem.2537.

Kumar P, Singh S, Handa V, et al. Oleic Acid Nanovesicles of Minoxidil for Enhanced Follicular Delivery. Medicines 2018;5(3):103; doi: 10.3390/medicines5030103.

Lorch M and Booth PJ. Insertion Kinetics of a Denatured α Helical Membrane Protein into Phospholipid Bilayer Vesicles. J Mol Biol 2004;344(4):1109–1121; doi: 10.1016/j.jmb.2004.09.090.

Mann S. The Origins of Life: Old Problems, New Chemistries. Angewandte Chemie International Edition 2013;52(1):155–162; doi: 10.1002/ANIE.201204968.

Martin W and Russell MJ. On the Origins of Cells: A Hypothesis for the Evolutionary Transitions from Abiotic Geochemistry to Chemoautotrophic Prokaryotes, and from Prokaryotes to Nucleated Cells. Philos Trans R Soc Lond B Biol Sci 2003;358(1429):59–85; doi: 10.1098/rstb.2002.1183.

Monnard P-A and Walde P. Current Ideas about Prebiological Compartmentalization. Life 2015;5(2):1239–1263; doi: 10.3390/life5021239.

Osaki T, Kuribayashi-Shigetomi K, Kawano R, et al. Uniformly-Sized Giant Liposome Formation with Gentle Hydration. In: 2011 IEEE 24th International Conference on Micro Electro Mechanical Systems IEEE; 2011; pp. 103–106; doi: 10.1109/MEMSYS.2011.5734372.

Rahimpour Y and Hamishehkar H. Liposomes in Cosmeceutics. Expert Opin Drug Deliv 2012;9(4):443–455; doi: 10.1517/17425247.2012.666968.

Sato W, Zajkowski T, Moser F, et al. Synthetic Cells in Biomedical Applications. WIREs Nanomedicine and Nanobiotechnology 2022;14(2):e1761; doi: 10.1002/wnan.1761.

Schrum JP, Zhu TF and Szostak JW. The Origins of Cellular Life. Cold Spring Harb Perspect Biol 2010;2(9):a002212–a002212; doi: 10.1101/cshperspect.a002212.

Smoukov SK, Seckbach J, Gordon R, et al. Origin of Life from a Maker’s Perspective–Focus on Protocellular Compartments in Bottom-Up Synthetic Biology. Conflicting Models for the Origin of Life 2023;303–326; doi: 10.1002/9781119555568.CH12.

Stano P, Rampioni G, Carrara P, et al. Semi-Synthetic Minimal Cells as a Tool for Biochemical ICT. Biosystems 2012;109(1):24–34; doi: 10.1016/J.BIOSYSTEMS.2012.01.002.

Subbotin V and Fiksel G. Exploring the Lipid World Hypothesis: A Novel Scenario of Self-Sustained Darwinian Evolution of the Liposomes. Astrobiology 2023;23(3):344–357; doi: 10.1089/ast.2021.0161.

Sun ZZ, Hayes CA, Shin J, et al. Protocols for Implementing an Escherichia Coli Based TX-TL Cell-Free Expression System for Synthetic Biology. Journal of Visualized Experiments 2013;(79); doi: 10.3791/50762.

Sur S, Fries AC, Kinzler KW, et al. Remote Loading of Preencapsulated Drugs into Stealth Liposomes. Proceedings of the National Academy of Sciences 2014;111(6):2283–2288; doi: 10.1073/pnas.1324135111.

van Swaay D and deMello A. Microfluidic Methods for Forming Liposomes. Lab Chip 2013;13(5):752–767; doi: 10.1039/c2lc41121k.

Tsumoto K, Matsuo H, Tomita M, et al. Efficient Formation of Giant Liposomes through the Gentle Hydration of Phosphatidylcholine Films Doped with Sugar. Colloids Surf B Biointerfaces 2009;68(1):98–105; doi: 10.1016/j.colsurfb.2008.09.023.

Venero OM, Sato W, Heili JM, et al. Liposome Preparation by 3D-Printed Microcapillary-Based Apparatus. Methods in Molecular Biology 2022;2433:227–235; doi: 10.1007/978-1-0716-1998-8_14.

Wang Q, Li W, Hu N, et al. Ion Concentration Effect (Na+ and Cl−) on Lipid Vesicle Formation. Colloids Surf B Biointerfaces 2017;155:287–293; doi: 10.1016/j.colsurfb.2017.04.030.

Zakir F, Vaidya B, Goyal AK, et al. Development and Characterization of Oleic Acid Vesicles for the Topical Delivery of Fluconazole. Drug Deliv 2010;17(4):238–248; doi: 10.3109/10717541003680981.

